# Task-Relevant Stimulus Design Improves P300-Based Brain-Computer Interfaces

**DOI:** 10.1101/2024.05.01.592004

**Authors:** Jongsu Kim, Yang Seok Cho, Sung-Phil Kim

**Affiliations:** Department of Biomedical Engineering, UNIST, Ulsan, Republic of Korea; School of Psychology, Korea University, Seoul, Republic of Korea

**Keywords:** P300-based BCI, Stimulus design, Task relevance, Selective attention, Single-trial BCI, Zero-calibration, Calibration-free, User Variability

## Abstract

**Objective:** In the pursuit of refining P300-based brain-computer interfaces (BCIs), our research aims to propose a novel stimulus design focused on selective attention and task relevance to address the challenges of P300-based BCIs, including the necessity of repetitive stimulus presentations, accuracy improvement, user variability, and calibration demands.

**Approach:** In the oddball task for P300-based BCIs, we develop a stimulus design involving task-relevant dynamic stimuli implemented as finger-tapping to enhance the elicitation and consistency of event-related potentials (ERPs). We further improve the performance of P300-based BCIs by optimizing ERP feature extraction and classification in offline analyses.

**Main Results:** With the proposed stimulus design, online P300-based BCIs in 37 healthy participants achieves the accuracy of 91.2% and the information transfer rate (ITR) of 28.37 bits/min with two stimulus repetitions. With optimized computational modeling in BCIs, our offline analyses reveal the possibility of single-trial execution, showcasing the accuracy of 91.7% and the ITR of 59.92 bits/min. Furthermore, our exploration into the feasibility of across-subject zero-calibration BCIs through offline analyses, where a BCI built on a dataset of 36 participants is directly applied to a left-out participant with no calibration, yields the accuracy of 94.23% and the ITR of 31.56 bits/min with two stimulus repetitions and the accuracy of 87.75% and the ITR of 52.61 bits/min with single-trial execution. When using the finger-tapping stimulus, the variability in performance among participants is the lowest, and a greater increase in performance is observed especially for those showing lower performance using the conventional color-changing stimulus.

**Signficance:** Using a novel task-relevant dynamic stimulus design, this study achieves one of the highest levels of P300-based BCI performance to date. This underscores the importance of coupling stimulus paradigms with computational methods for improving P300-based BCIs.

## 1. Introduction

Brain-computer interfaces (BCIs) are rapidly evolving at the intersection of neuroscience and engineering, offering revolutionary communication and control channels independent of peripheral neural and muscular activity [1]. A P300-based BCI is one of the non-invasive BCIs that leverages the P300 component in event-related potentials (ERPs) of electroencephalography (EEG), which is elicited by recognizing a target stimulus in the oddball paradigm[2]. P300-based BCIs have been widely adopted in various applications from assistive technologies to gaming and virtual reality [3, 4]. However, P300-based BCIs still face challenges, including repeated stimulus presentations, accuracy enhancement, individual variation, and frequent calibration requirements, which need to be addressed through innovative approaches for the enhancement of usability and applicability.

### 1.1. Challenges in P300-based BCIs

First, P300-based BCIs generally require repeated stimulus presentations for accurate ERP detection, limiting real-world application efficiency [5-9]. Despite advancements in single-trial P300-based BCIs, their performance falls short compared to multi-trial approaches [10, 11]. Second, P300-based BCIs necessitate personalized feature extraction and decoding algorithms as well as separate calibration sessions for each use, thus hindering daily practicality [12]. Transfer learning strategies have been developed to minimize the extensive calibration requirements; yet, attaining the robustness necessary for consistent applications across diverse datasets has proven to be challenging [13-16]. Third, the P300 component depends on cognitive states such as attention and working memory. Thus, strategies such as increasing stimulus saliency are necessary to enhance the quality of ERPs [4, 17-19]. These challenges emphasize the need for balanced P300-based BCI designs that can effectively integrate technical as well as user-centered aspects.

### 1.2. System Design of P300-based BCIs

To address the challenges in P300-based BCIs above, we need to improve the design of both computational methods and task paradigms. Computational methods can be designed using advanced signal processing and machine learning techniques to enhance decoding performance. For instance, the number of stimulus presentations can be reduced by adaptively determining an optimal point to halt the stimulation process, thereby enhancing information transfer rates (ITRs) while maintaining accuracy [20]. Enlarging training datasets with transfer learning or generative artificial intelligence (AI) methods can also improve decoding models in P300-based BCIs [21].

While adopting advanced computational methods has the potential to provide a promising solution to the challenges in P300-based BCIs, improving decoding models alone may require extensive exploration of optimal algorithms and enormous efforts to validate their efficacy across diverse BCI applications. For instance, variations in the P300-based BCI performance across users partially due to ‘BCI illiteracy’ remain a problem that has not been completely resolved by decoding improvement [19]. While decoding models are a key component of the BCI system, another component to elicit high-quality ERPs via innovative task paradigms is also critical. In line with the principle of ‘garbage in, garbage out’, high-quality ERPs would lead to robust BCI performance.

As such, we assume that the challenges in P300-based BCIs can be addressed by not only advanced computational models but also effective paradigm designs. Specifically, an effective paradigm design for P300-based BCIs may enhance the elicitation of ERP responses in the oddball task. Paradigm design factors in the oddball task generally include target probability, inter-stimulus interval (ISI), inter-target interval (ITI), and stimulus repetition, which affect the P300 amplitude and waveform [4, 8, 22, 23]. Yet, in this study, we focus on selective attention which plays a key role in cognitive processing during the oddball task [4]. We hypothesize that if we can elevate selective attention to target stimuli in the oddball task, the corresponding ERPs would be more reliably elicited across individuals with fewer stimulus repetitions.

### 1.3. Attention Considerations in P300-based BCI Design

To elevate selective attention in the oddball task, we design stimuli by considering several attentional aspects: bottom-up attention, top-down attention, and task relevance [24, 25]. From a bottom-up perspective, many studies have suggested various stimulus designs to attract user attention to induce more pronounced ERPs [3, 12, 18, 26-31]. In particular, we intend to make stimuli dynamic as ERP studies indicate that using dynamic stimuli—moving stimuli that evoke motion-onset visual evoked potentials (mVEPs) [28-30]— is a strategic choice to enhance the natural “attention-grabbing” capacity and reduce intra-subject variability in ERPs [28].

Top-down attention is also crucial. P300-based BCI studies have explored designs for top-down attention with physical responses (e.g., button pressing) or mental tasks (e.g., counting) [32] or by considering a relationship between task difficulty and P300 amplitude [4, 22, 33]. In particular, mental tasks are more viable in practical BCI use as they entail active engagement, reportedly producing larger ERP responses than merely paying attention to stimuli [28, 34, 35]. However, other mental tasks than counting have been scarcely employed for P300-based BCIs, in contrast to motor imagery (MI)-based BCIs that have explored diverse mental tasks [36, 37].

Contrary to bottom-up and top-down attention, task relevance has not yet been taken into account when designing P300-based BCIs. While motor imagery (MI)-BCIs have extensively explored the task-relevance aspects [23, 24], their investigation in reactive BCIs, particularly P300-based BCIs has been limited [25, 26]. This disparity possibly arises because active BCIs, such as MI-BCIs, require users to generate brain signals through imagination, whereas reactive BCIs, such as P300-based BCIs, rely on stimuli-induced signals, leading research to focus primarily on attention mechanisms. However, studies indicate that the presentation of stimuli that are more relevant to tasks can enhance ERP responses [38]. As such, we assume that designing stimuli that are relevant to the oddball task would help to enhance ERPs in P300-based BCIs. As the oddball task involves selective attention, we aim to design stimuli related to the task of “selectively attending to a target”.

Considering the divserse aspects of attention mentioned above, we design a finger-tapping stimulus inspired by the fact that people frequently use finger-tapping to select items on touch screens in their daily lives. Therefore, the finger-tapping stimulus would be more relevant to the oddball task than conventional stimuli. The finger-tapping stimulus is also designed to guide a mental task, other than counting, to imagine finger movements. We investigate whether such an MI task would improve ERPs compared to the counting task. Lastly, we make the finger-tapping stimulus dynamic (i.e., animated) to increase saliency in bottom-up attention.

We aim to investigate whether the proposed paradigm centered on selective attention and task relevance can minimize repetition needs and improve ERP consistency across individuals. We expect that such improved ERP consistency would facilitate transfer learning across users [39, 40]. In addition, the proposed paradigm using dynamic stimuli to enhance bottom-up attention would further reduce the variability of BCI performance across users [28].

### 1.4. Optimization of P300-based BCI design

With a novel stimulus design proposed in this study, we aim to optimize the design of P300-based BCIs by incorporating paradigm design and computational methods together. As the proposed stimulus design is expected to generate more reliable ERPs, we need to investigate proper computational methods to decode them. So, we explore various signal processing and decoding models that best fit ERPs elicited by the finger-tapping stimulus. By combining the resultant computational methods with the finger-tapping stimulus, we aim to address the aforementioned challenges in P300-based BCIs, including stimulus repetitions, individual variation, and calibration requirements.

To assess our novel design for P300-based BCIs, we first conduct an online BCI experiment to evaluate the effect of the finger-tapping stimulus design on the BCI performance. In this experiment, we solely focus on the paradigm design without optimizing the design of computational models. We compare the online BCI performance between the design of the finger-tapping stimulus with the MI task and other stimulus designs controlling each aspect of attention: bottom-up (dynamic vs. static), top-down (MI vs. counting task), and task relevance (relevant vs. irrelevant to selection). Here, we use conventional computational models to highlight the effect of paradigm designs.

Next, we optimize the computational models for the finger-tapping stimulus design in a separate offline analysis. Based on ERPs elicited by the finger-tapping stimulus, we seek to determine computational models to minimize stimulus repetitions (towards single-trial), remove individual calibration (zero-calibration), and reduce variations in performance across subjects,

## 2. Methods

### 2.1. Participants

Based on the statistical power analysis using G*Power 3 [41], a sample size of 33 participants was determined for the experiment (effect size f = 0.25, power = 0.95, nonsphericity correction ε = 0.8). To meet this minimum requirement, we recruited 37 healthy adults (21.62±3.53 years old, range: 18-30 years old, 8 females). All participants were right-handed and reported no history of mental illness or neurological disorders. Informed consent was obtained from participants in compliance with the Ulsan National Institutes of Science and Technology, Institutional Review Board (UNIST-IRB-18-08-A).

### 2.2. Stimulus Design

We crafted visual stimuli that were tailored by our previous work for the integration of BCIs into different user interfaces (UIs) across monitors, augmented reality (AR), and virtual reality (VR) environments [12]. Positioned at the four quadrants of the display (top-left, bottom-left, top-right, bottom-right), these stimuli featured icons indicative of on/off, play, stop, and pause functions, set against a blue rectangular background, mirroring the potential control of an external device (i.e., Bluetooth speaker in this study) [12].

We created three types of stimuli. First, a static stimulus was crafted that changed the color of the rectangular background from blue to green (Figure 1(a)). This control stimulus was supposed to be the least salient in bottom-up attention among other stimuli used in this study and task-irrelevant. Second, a dynamic stimulus was built by incorporating animations on top of the color-changing stimulation (Figure 1(b)). The stimulus was animated by rotating the icons (i.e., rotating the icons by 90° clockwise), similar to other studies [42]. This second stimulus was supposed to be more salient than the first one but still task-irrelevant. Note that we added animation to color-changing effects when designing the second stimulus since we intended to ensure that its bottom-up saliency was greater than the first one. Third, we created the finger-tapping stimulus, designed to be task-relevant and dynamic, by animating the right index finger to press the icons and return to an initial position (Figure 1(c)). Again, the finger-tapping animation was added to the color-changing stimulation to increase bottom-up saliency. The third stimulus was distinct from the second one by making the animation task relevant.

**Figure 1.**
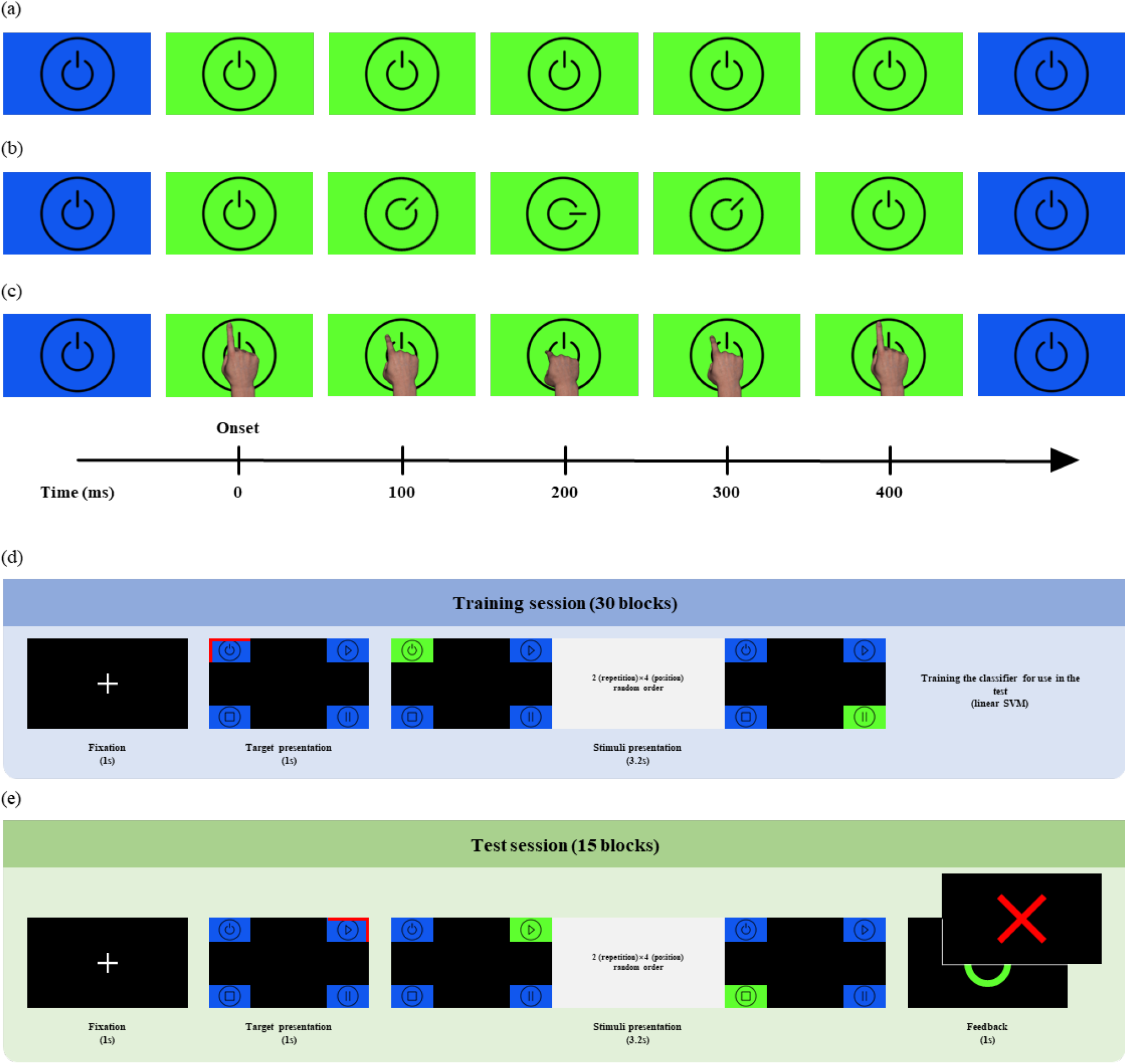
Stimulus designs and online experiment procedure. An example control option for a Bluetooth speaker, turning on the speaker, is displayed with three different visual stimulation designs: (a) Static color-changing stimulus, (b) Task-irrelevant dynamic icon-rotating stimulus, and (c) Task-relevant dynamic finger-tapping stimulus. Each stimulus is presented for 400 ms. The online experiment undergoes the training and test sessions. (d) The training session include 30 blocks, each of which begins with a 1-second fixation, followed by 1-second target presentation and 3.2-second stimulus presentation where each of four stimuli is presented twice sequentially in a random order at four corners. After all blocks, a linear SVM classifier is trained using the training session data. (e) The test session includes 15 blocks mirroring the training session structure. A difference is that after stimulus presentation, BCI control feedback is given to participants for 1 second, indicsting whether the target is accurately decoded.

Each stimulus was presented for 400 ms. This stimulus duration (SD) was determined to render the appearance of finger-tapping as natural as possible within the shortest time frame. Given the importance of the total stimulus presentation duration for practical use in P300-based BCIs, we fixed the ISI to 0 ms.

### 2.3. Task Design

In the experimental setup, participants performed two specific mental tasks when they saw a target stimulus: counting and MI. In the counting task, participants simply counted the number of times the target stimulus appeared. In the MI task, participants were instructed to imagine the movement of flexion and extension of the right index finger. To account for differences in individual performance of MI, a guideline was provided using visual examples of finger-tapping that matched the stimulus design. Participants were then asked to mentally replicate these finger movements.

### 2.4. Online Experimental Procedure

Throughout the online experiment of P300-based BCIs, participants were comfortably seated in front of a 27-inch monitor. Before the start of the experiment, participants read a manual detailing the experimental procedure, received thorough explanations, and had a question-and-answer session. To ensure the successful performance of MI during the experiment, sufficient training was provided with the guideline described above until participants felt confident to proceed with their MI. They were instructed to self-control the progression of the experiment by pressing the space bar (see below for details). An electromyography (EMG) electrode was placed over each of the metacarpophalangeal and distal interphalangeal joints of the participant’s right index finger to monitor potential muscle activity during the MI. Before the experiment began, we confirmed that deviant EMG was observed when participants flexed or extended their right index finger.

A block of the P300-based BCI operation underwent four phases as follows (Figure 1(d, e)): 1) Fixation: participants gazed at a central cross for 1,000 ms; 2) Target indication: the target location among four corners was indicated by displaying a red border around the corresponding icon for 1,000 ms; 3) Stimulus presentation: each icon was sequentially changed twice according to a given stimulus type in a randomized order, resulting in 2 repetitions of stimulus presentation; and 4) Feedback presentation (only for test blocks): participants received immediate feedback on whether the BCI system correctly identified the target stimulus, where the BCI output was displayed on the screen for 1,000 ms. Participants were instructed to pay attention to the presentation of a target stimulus while performing a given mental task. They were given 1 target and 3 non-target stimuli twice for a total of 3,200 ms (8×400 ms). During training, the feedback presentation phase was omitted. Each stimulus presentation will be referred to as a trial hereafter.

The experiment consisted of 6 sessions of blocks by combining 3 stimulus types with 2 mental tasks: color changing with counting, color changing with MI, icon-rotating with counting, icon-rotating with MI, finger-tapping with counting, and finger-tapping with MI. The order of 6 sessions was randomized for each participant. Participants performed 45 blocks per session, with 30 blocks of training and 15 blocks of testing.

Before the main experiment, participants were engaged in 6 practice sessions, each consisting of five blocks, to become acquainted with the tasks and stimuli. The practice sessions were designed in the following sequence: 1) color changing with counting, to introduce participants to a basic P300-based BCI task; 2) finger-tapping with counting, and 3) finger-tapping with MI, to help participants learn to synchronize their MI with the finger-tapping stimulus; 4) color changing with MI, to continue practicing MI with a different stimulus, reinforcing what was learned in the previous sessions 1); and finally, 5) icon-rotating with counting and 6) icon-rotating with MI, to expose participants to novel stimuli and practice MI, thereby increasing their adaptability to various BCI tasks. With this structured practice, we intended participants to be prepared for different tasks in the main experiment.

### 2.5. Data Acquisition and Preprocessing

EEG data were recorded using 31 active wet electrodes (FP1, FPz, FP2, F7, F3, Fz, F4, F8, FT9, FC5, FC1, FC2, FC6, FT10, T7, C3, Cz, C4, T8, CP5, CP1, CP2, CP6, P7, P3, Pz, P4, P8, O1, Oz, and O2) following the international 10–20 system (American Clinical Neurophysiology Society Guideline 2). Signals were transmitted to an EEG amplifier (actiCHamp, Brain Product GmbH, Gilching, Germany) at a 500-Hz sampling rate. The left and right mastoids were used as reference and ground, respectively. Electrode impedance was kept below 10kΩ throughout the experiment.

EEG preprocessing underwent in the following order: (1) 1-Hz high-pass filtering, (2) 50-Hz low-pass filtering, (3) bad-channel removal and interpolation, (4) re-referencing using common average reference (CAR), (5) artifact removal using artifact subspace reconstruction (ASR) (cutoff 10), and (6) 12-Hz low-pass filtering. Filters were designed as finite impulse response (FIR) filters using a Hamming window to attenuate unwanted frequencies. Bad channels were detected by using a modified version of the ‘clean_channels’ function from the EEGLAB toolbox. A channel’s correlation with its neighbors was evaluated over 5 seconds, with those showing a correlation coefficient below 0.8 deemed suspect. Channels were classified as ‘bad’ if their abnormal state persisted over 40% of the recordings, ensuring the exclusion of consistently noisy data from subsequent analyses.

EMG signals were recorded with 2 passive wet electrodes at metacarpophalangeal and distal interphalangeal joints of the participant’s right index finger. EMG signals were transmitted to an EEG amplifier through a BIP2AUX adapter (Brain Products GmbH, Gilching, Germany), allowing for integration and recording at the same sampling rate as the EEG data. EMG preprocessing was conducted as follows: (1) 1-Hz high-pass filtering to remove low-frequency drifts, (2) 150-Hz low-pass filtering to eliminate high-frequency noise, and (3) 60-Hz notch filtering to attenuate power line interference. Subsequently, the signals were rectified, and smoothed using a 50-ms moving average.

In the online experiment, preprocessing was conducted after the completion of all 30 blocks of the training part. For the test part, which consisted of 15 blocks, preprocessing was performed immediately after each block. Due to the real-time operation of BCIs during testing, re-applying bad-channel removal, interpolation and ASR could be problematic in the preprocessing of the test block data. Thus, we adopted the bad-channel information and ASR parameters obtained from the preprocessing of the training data, instead of recalculating them for each test block.

### 2.6. Online BCI Decoding

EEG signals at each channel were epoched from -200 ms to 600 ms relative to the stimulus onset. The epoched EEG signals were baseline-corrected by subtracting the mean amplitude during the pre-stimulus period. Averaging these signals over the two trials resulted in an ERP waveform for each stimulus.

ERP features were extracted simply by concatenating the ERP waveforms within a post-stimulus period (0 ∼ 600 ms) from all channels into a 1D vector that contained 9,300 features (31 channels × 300 time points) for each stimulus. A linear support vector machine (SVM) classifier was trained using 120 feature vectors from the 30 training blocks; 30 target stimuli labeled as +1 and 90 non-target stimuli labeled as -1. Then, in each testing block, we created feature vectors in the same way as in training, yielding 4 vectors corresponding to each stimulus. The trained linear SVM classifier produced the classification score for the *j*-th stimulus as follows:

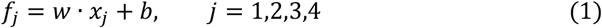

where *w* is the weight vector consisiting of support vectors in linear SVM, *x*_*j*_ is the feature vector for the *j*-th stimulus, and *b* is a bias term. The stimulus with the highest score (*f*_*j*_) was predicted as a target.

### 2.7. BCI Performance Evaluation

The performance of the BCI system was quantified using three metrics: accuracy, ITR, and coefficient of variation (CV).

Accuracy was defined as the ratio of the number of successful test blocks to the total number of test blocks. A test block was successful if the BCI system correctly identified a test stimulus.

ITR was calculated in bits per minute (bits/min) using the following formula:

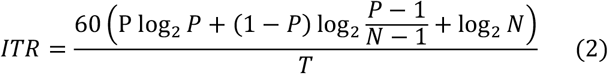

where *P* is accuracy, *N* is the number of selectable options (*N*=4 here), and *T* is the time taken for one selection in seconds (*T*=(SD+ISI)×*N*×repetition of stimuli).

CV was used to assess the consistency of accuracy and ITR among participants, calculated as the standard deviation divided by the mean of accuracy (or ITR) across participants.

### 2.8. Offline Optimization of Computational Models

In the post-hoc offline analysis of the online BCI experimental data, we optimized the design of computational models to further improve P300-based BCIs. The optimization was conducted to tackle the aforementioned three challenges in P300-based BCIs: the need for multiple stimulus repetitions, individual calibration requirements, and individual variations in performance. The optimization of computational models was undertaken in two parts: feature extraction and classification.

Among many models to extract ERP features, we opted for spatial filtering and manifold transformation. First, we chose to use xDAWN for spatial filtering as it is known to extract ERP features appropriately in a supervised manner, particularly the P300 component [43]. The ability of xDAWN to enhance ERP features was expected to mitigate the limited quality of ERPs associated with reduced stimulus repetitions. Second, we chose to use Riemannian geometry (RG) to transform ERP features onto manifolds as it is adept at representing complex data structures and well-suited for addressing EEG signal complexity. Especially, using xDAWN and RG together has proven to enhance P300-based BCI performance [14, 44]. For feature extraction, we first generated ERPs by utilizing the post-stimulus period of the baseline-corrected signals, as described in Section 2.6. We then applied xDAWN spatial filtering to these ERPs and calculated the symmetric positive definite (SPD) covariance matrices of the filtered signals. These SPD matrices were represented on the Riemannian manifold and projected onto the tangent space to transform ERP features onto manifolds [44]. We utilized the pyRiemann Python package [45] for xDAWN filtering and RG computation.

As for classification models, we investigated both conventional machine learning models and deep learning models that have been used for BCIs. Linear SVM and logistic regression (LR) were adopted as conventional machine learning models. For deep learning, we used EEGNet [46], shallow ConvNet [47], and deep ConvNet [47], which have been particularly shown to be suitable for classifying P300-based BCI data.

We optimized computational models for P300-based BCIs in a greedy manner by examining the best combination of feature extraction and classification models: 3 feature extraction models (none, xDAWN (XD), xDAWN, and RG (XDRG)) and 5 classification models (SVM, LR, EEGNet, shallow ConvNet, and deep ConvNet). We sought the best combination for individual problems as described below.

#### 2.8.1. Minimization of Stimulus Repetitions

Although we already reduced the number of stimulation presentations to 2 in our online experiment, we investigated if we could further reduce it to 1 to realize a single-trial P300-based BCI (i.e., no repetition). To this end, we selected EEG data in the first round of stimulus presentation from the online experiment and explored a combination of feature extraction and classification models that produced the highest BCI performance when stimuli were presented once. For comparison, we also optimized feature extraction and classification models offline when stimuli were presented twice.

#### 2.8.2. Across-Subject Zero-Calibration

In our pursuit of a zero-calibration, plug-and-play P300-based BCI, we applied transfer learning to the online experiment data with the Leave-One-Subject-Out Cross-Validation (LOSO CV) scheme. The training set comprised all 45 blocks from all subjects except one, whose testing blocks (15 blocks) were used for validation. We explored the best combination of feature extraction and classification models. Furthermore, as in section 2.8.1, by analyzing first-trial data, we also explore the possibility of achieving zero-calibration at the single-trial level. This cross-validation demonstrates the feasibility of zero-calibration that can realize a plug-and-play P300-based BCI.

#### 2.8.3. Individual Variation of BCI Performance

We explored whether the proposed BCI design could reduce individual variations of BCI performance. To this end, we analyzed the CV values across 6 different BCI paradigms. Furthermore, we assessed whether the proposed design could improve more the performance of participants who showed relatively lower performance using conventional BCI designs – i.e., the color-changing stimulus with counting in our case. We first evaluated changes in BCI performance from using the color-changing stimulus with counting to using the finger-tapping stimulus with counting. Then, we calculated Pearson’s correlation coefficient between the performance using the color-changing design and the performance change induced by the finger-tapping design.

### 2.9. Statistical Analysis of BCI Performance

A 2-way repeated-measures ANOVA (rmANOVA) was employed to assess the BCI performance across different paradigms. The analysis focused on two dependent variables: accuracy and ITR. The independent variables under examination were the stimulus type (with 3 levels: static as in color change, and dynamic as in icon-rotating and finger tapping) and the mental task (with two levels: counting and MI of finger tapping). To assess the statistical differences in performance across various combinations of sessions and analysis methods, a one-way repeated-measures ANOVA (one-way rmANOVA) was employed. This statistical approach was particularly utilized to analyze differences among distinct scenarios combining different sessions and computational strategies. When the assumption of sphericity was violated, the Greenhouse-Geisser correction was applied to ensure the validity of the rmANOVA results. Post hoc analyses, using Tukey’s Honestly Significant Difference Procedure (HSD), further dissected the effects of stimulus types, mental tasks, and their interaction on BCI performance. For the offline analysis, we utilized a paired t-test to compare the efficacy between classification methods.

## 3. Results

### 3.1. Online BCI Control

We assessed the performance of online P300-based BCIs with 6 different designs in terms of accuracy and ITR. We found the highest accuracy with the finger-tapping stimulus and counting, followed by the finger-tapping stimulus and MI, the icon-rotating and MI, the icon-rotating and counting, the color-changing and MI, and the color-changing and counting (see Table 1 for details). The highest accuracy with the finger-tapping and counting was 91.17% on average whereas the lowest with the color-changing and counting was 80.00%, resulting in the improvement of 11.17% by adopting a new BCI paradigm (Figure 2(a)). Similar trends were observed in ITR, where using the finger-tapping stimuli outperformed other paradigms (Figure 2(b)).

**Table 1.** Online P300-Based BCI performance assessed by accuracy and ITR for three stimulus designs combined with two mental tasks. Mean and standard deviation across N=37 participants. Bold fonts represent the highest performance. # stimulus repetitions = 2.

| Stimulus designs | Mental task | Accuracy |  | ITR (bits/min) |  |
| --- | --- | --- | --- | --- | --- |
|  |  | Mean | Std. | Mean | Std. |
| Color-changing | Counting | 0.8000 | 0.1663 | 20.3615 | 10.2798 |
|  | MI | 0.8018 | 0.1760 | 20.8218 | 11.0490 |
| Icon-rotating | Counting | 0.8541 | 0.1285 | 23.7372 | 9.4733 |
|  | MI | 0.8631 | 0.1227 | 24.5625 | 9.8569 |
| Finger-tapping | Counting | **0.9117** | 0.0917 | **28.3704** | 8.4638 |
|  | MI | 0.8955 | 0.0988 | 26.8445 | 8.4952 |

**Figure 2.**
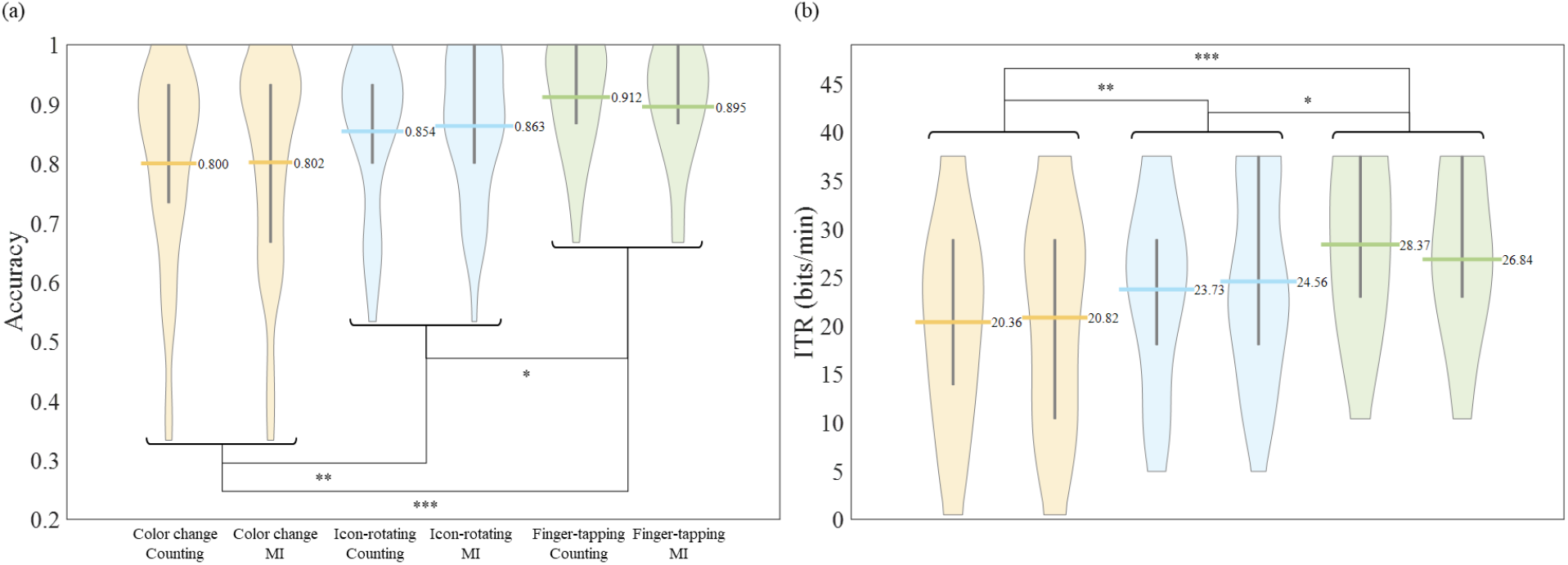
Online P300-based BCI performance with six different paradigms, including color-changing, icon-rotating and finger tapping stimulus designs combined with the counting and motor imagery tasks. (a) Distributions of accuracy across N=37 participants. (b) Distributions of information transfer rate (ITR). The bold horizontal line represents the mean, and the ends of the gray vertical lines denote the lower and upper quartiles. Statistical significance is denoted as follows: *p < 0.05, **p < 0.01, ***p < 0.001, Tukey’s HSD post-hoc test.

A 2-way rmANOVA on accuracy revealed the significant main effect of the stimulus type (p =3.4834×10^-9^), while neither the main effect of the mental task nor the interaction effect was found. A post-hoc analysis showed significant differences between color-changing and finger-tapping (p =7.8790×10^-6^), color-changing and icon-rotating (p = 0.0037), and finger-tapping and icon-rotating (p = 0.0279), affirming the superior performance by using the finger-tapping stimulus. Similary, a 2-way rmANOVA on ITR showed the significant main effect of the stimulus type (p = 5.3242×10^-7^). A post-hoc analysis also showed significant differences between the stimilus types: color-chaning < icon-rotating < finger-tapping (ps < 0.05). We verified that the EMG amplitude was not different between the MI and counting tasks for each of the three stimulus designs (paired t-test, ps > 0.05), which indicates that MI did not modulate the EMG signals in the experiment.

### 3.2. Optimization toward Single-Trial BCI

In the offline analysis of BCI performance depending on the number of stimulus presentation repetitions, among all combinations of 3 feature extraction methods (none, XD, XDRG) and 5 classifiers (SVM, LR, EEGNet, shallow ConvNet, and deep ConvNet), using XDRG and LR produced the highest accuracy and ITR when we used single-trial ERPs as well as when we used the average ERPs from 2 repetitions (see Table S1 for the full results of all combinations). The highest performance was achieved with the finger-tapping stimlus and counting, echoing the findings from Section 3.1.

Using the paradigm of the finger-tapping stimulus with counting and the optimized models (XDRG and LR), one-way rmANOVA revealed a significance difference in accuracy among three cases: online 2 repetitions without optimization, offline 2 repetitions with optimization, and offline no repetition with optimization (p = 0.0059). Offline accuracy from 2 repetitions of stimulus presentation with optimization was higher than offline accuracy (Figure 3(a)) from single-trial presentation with optimization (p < 0.05) and online accuracy from 2 repetitions of stimulus presentation without optimization (p < 0.001). However, there was no significant difference in accuracy between offline single-trial presentation with optimization and online 2 repetitions of presentation without optimization. One-way rmANOVA also revealed a significance difference in ITR among three cases. ITR was the highest from offline single-trial presentation compared to offline and online ITRs from 2 repetitions of presentation (Figure 3(b)), due to reduced presentation duration (p < 0.001). Therefore, the offline performance of single-trial P300-based BCIs with optimized computational models yielded a similar level of accuracy and doubled improvement of ITR compared to the online performance of mult-trial (2 repetitions) P300-based BCIs without optimized computational models (see Table 2).

**Table 2.**
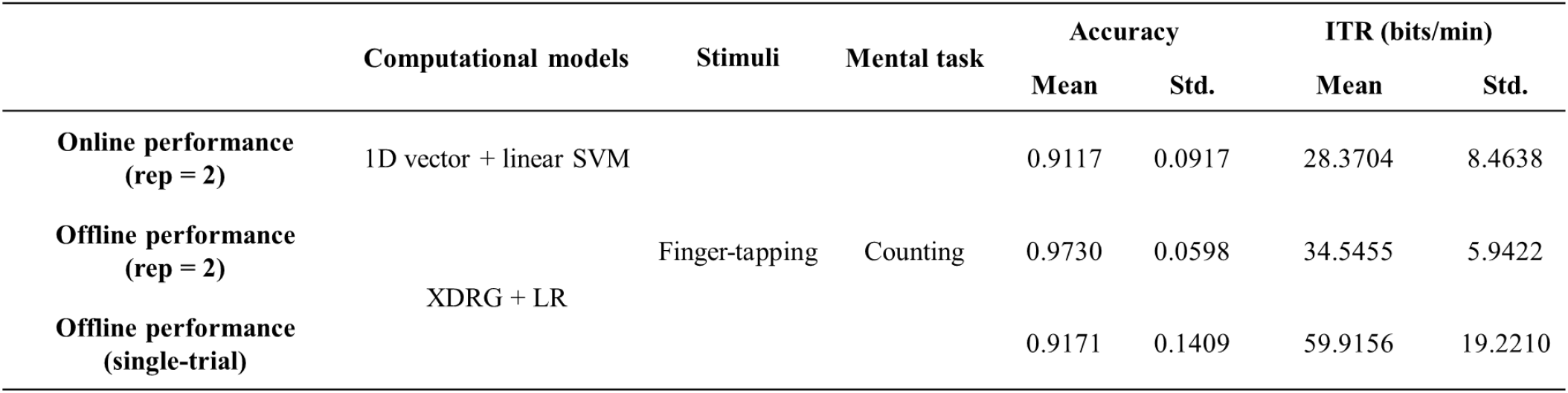
The performance of three P300-based BCIs, including online performance with 2 stimulus repetitions (rep=2), offline performance with 2 stimulus repetitions (rep=2) and computational models optimized, and offline performance with single-trial presentation and computational models optimized. Refer to Figure 3.

**Figure 3.**
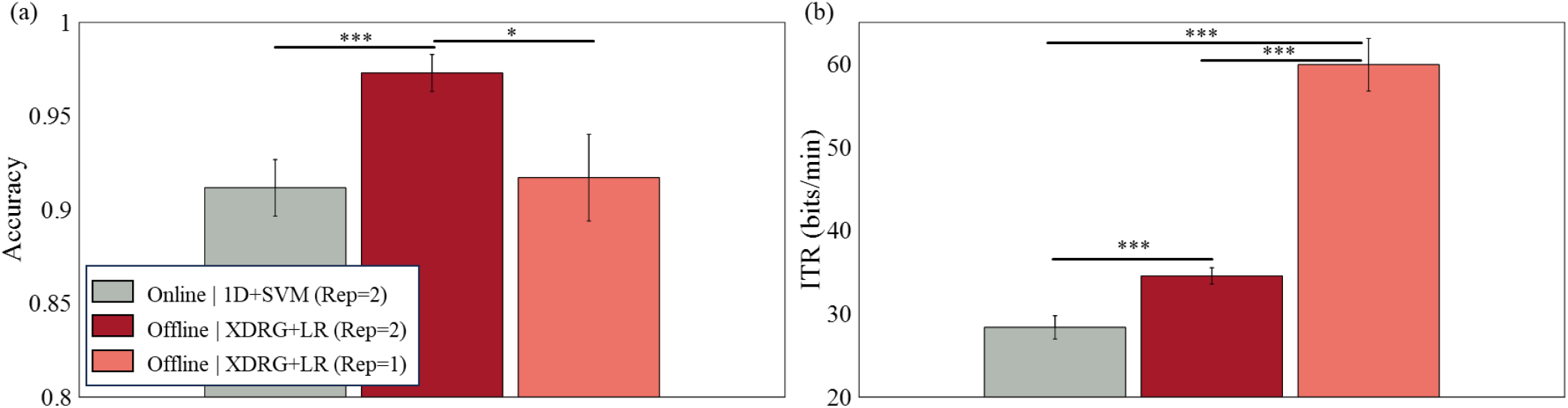
The performance evaluation of three P300-based BCIs: 1) online performance with 2 stimulus repetitions (Rep = 2); 2) offline performance with 2 stimulus repetitions (Rep = 2) and computational models optimized; and 3) offline performance with single-trial (Rep=1) and computational models optimized. The stimulus design of finger-tapping with counting task is evaluated. (a) Accuracy. (b) ITR. The feature extraction method labelled ‘1D’ concatenates ERPs of each channel into a 1D vector and that labelled ‘XDRG’ applies xDAWN spatial filtering and projects onto a Riemannian geometry manifold. Classification models are either a linear Support Vector Machine (SVM) or logistic regression (LR). Statistical significance is indicated as *p < 0.05, ***p < 0.001. Tukey’s HSD post-hoc test. Error bar denotes standard error of mean (SEM) (N = 37).

### 3.3. Optimization toward Zero-Calibration

In the offline analysis of zero-calibration BCIs via transfer learning, among all combinations of 3 feature extraction methods and 5 classifiers, using XD and deepConvNet produced the highest accuracy and ITR from LOSO cross-validation (see Table S2 for full results of all combinations). Again, the highest zero-calibration BCI performance was achieved using the finger-tapping stimulus with counting (see Table 3).

**Table 3.**
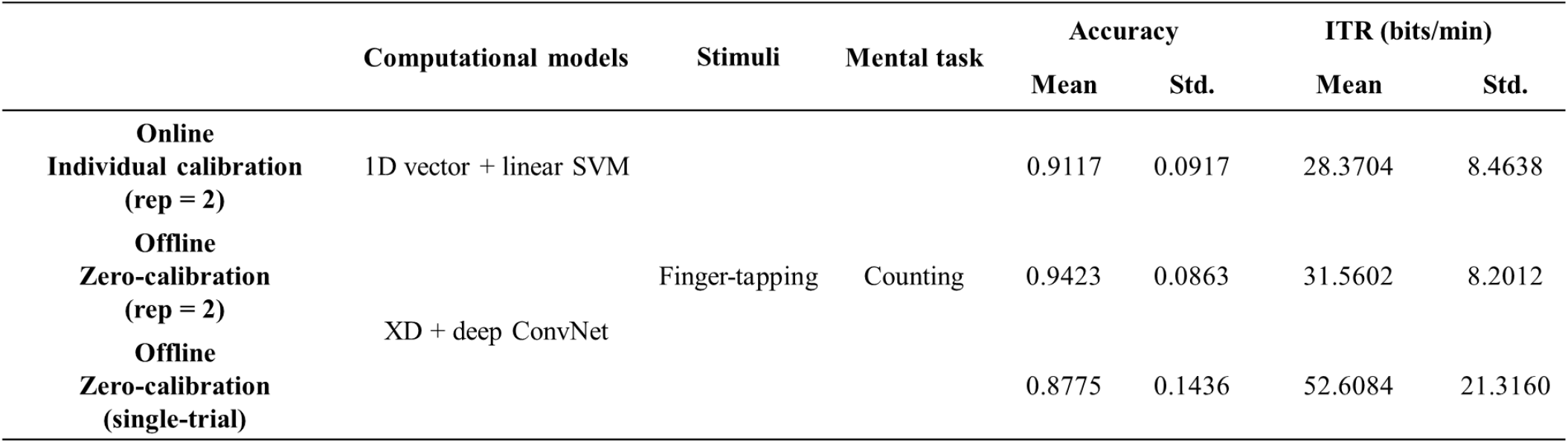
Performance of three P300-based BCIs: 1) online performance with 2 stimulus repetitions (rep=2) and individual calibration; 2) offline performance with 2 stimulus repetitions (rep=2), computational models optimized, and zero-calibration; and 3) offline performance with single-trial, computational models optimized, and zero-calibration. Refer to Figure 4.

Using the finger-tapping stimulus with counting and optimized models (XD and deepConvNet), one-way rmANOVA showed a significant difference in accuracy among three cases: online individual calibration of 2 repetitions without optimization, offline zero-calibration of 2 repetitions with optimization, and offline zero-calibration of no repetition with optimization (p= 0.0038) (Figure 4(a)). A post hoc analysis showed that the accuracy in offline zero-calibration with two stimulus repetitions surpassed that of no calibration (p < 0.001). There was no difference in accuracy between online individual cabliration without optimization and offline zero-calibration with optimization in 2 repetitions (p = 0.2122) and no repetition (p = 2183). Also, one-way rmANOVA showed a significant difference in ITR among three cases (p = 1.6628×10^-11^) (Figure 4(b)). The hightest ITR was achieved in offline single-trial presentations, outperforming both offline and online ITRs with two repetitions (p < 0.001). There was no significnat differencde in ITR between online individual calibration without optimization and offline zero-calibration with optimization for two repetitions of stimulus presentation. These results, consistent with those in Section 3.2., indicate the potential for both single-trial and zero-calibration P300-based BCIs.

**Figure 4.**
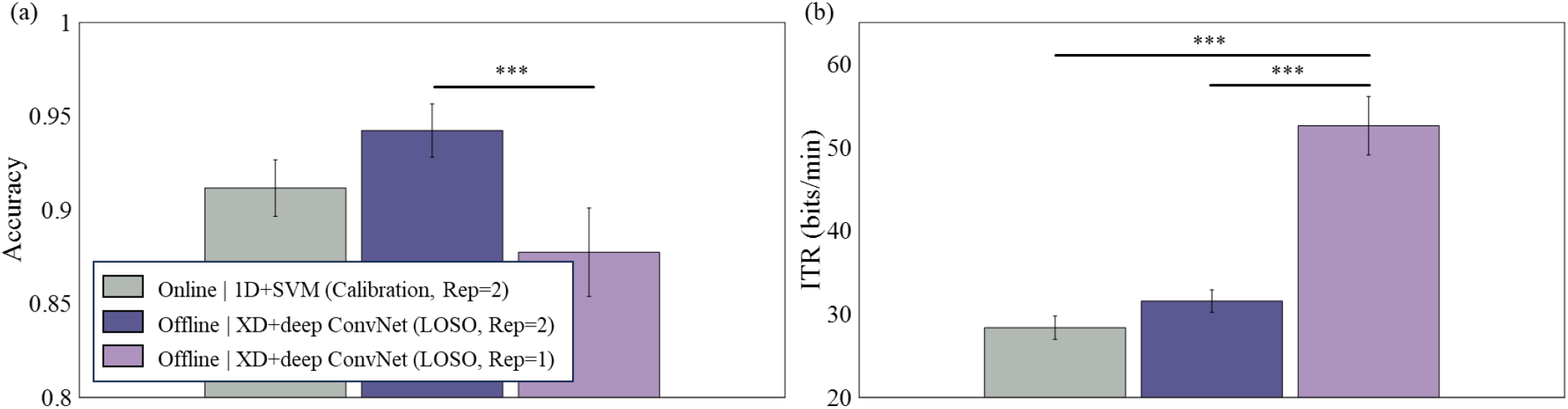
The performance evaluation of three P300-based BCIs: 1) online performance with 2 stimulus repetitions (Rep=2) and individual calibration; 2) offline performance with 2 stimulus repetitions (Rep=2), computational models optimized, and zero-calibration; and 3) offline performance with single-trial (Rep=1), computational models optimized, and zero-calibration. The stimulus design of finger-tapping with counting task is evaluated. (a) Accuracy. (b) ITR. The feature extraction method labelled ‘1D’ concatenates ERPs of each channel into a 1D vector and that labelled ‘XD’ applies xDAWN spatial filtering. Classification models are either a linear Support Vector Machine (SVM) or deep ConvNet. Zero-calibration is evaluated through leave-one-subject-out (LOSO) scheme. Statistical significance is indicated as ***p < 0.001. Tukey’s HSD post-hoc test. Error bar denotes standard error of mean (SEM) (N=37).

### 3.4. Individual Variation of BCI Performance

The analysis of CV revealed a trend that using dynamic stimuli exhibited lower CV values compared to using static stimuli (Figure 5). Moreover, using the finger-tapping stimuli reduced CV more than using other stimuli for all online and offline analyses (see Sections 3.2 and 3.3 for details of each analysis). The lowest CV was observed in the XDRG with LR during the offline analysis of BCIs with 2 repetitions of finger-tapping stimuli. In this case, the accuracy ranged from 0.7333 to 1 and the ITR ranged from 13.8882 to 37.5 bits/min across 37 participants.

**Figure 5.**
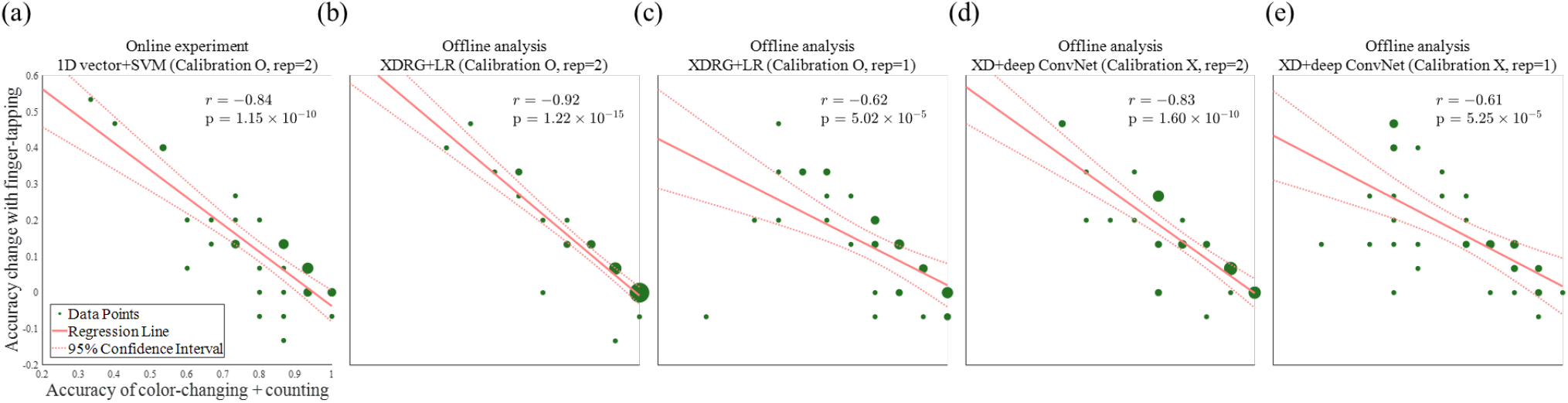
Across-subject correlations between P300-based BCI accuracy using the color-changing stimulus and changes in P300-based BCI accuracy using the finger-tapping stimulus relative to that using the color-changing stimulus. The mental tasks is counting. The solid red line represents linear regression estimation, while the red dashed lines indicate 95% confidence intervals. Each green dot represents individual participants, with larger dots indicating multiple participants sharing the same value. r denotes correlation coefficient with the p-value (p) resulted from the F-test. Correlations are evaluated for five different P300-based BCI setups (see Figure 5 for details).

Additionally, significant correlations were observed across all online and offline analyses between the accuracy in the color-changing with counting sessions and the change in accuracy when using finger-tapping stimuli (ps < 0.05) (Figure 6). Negative correlations showed that participants having lower accuracy in the color-changing with counting tended to achieved more improvement in accuracy by the finger-tapping with counting.

**Figure 6.**
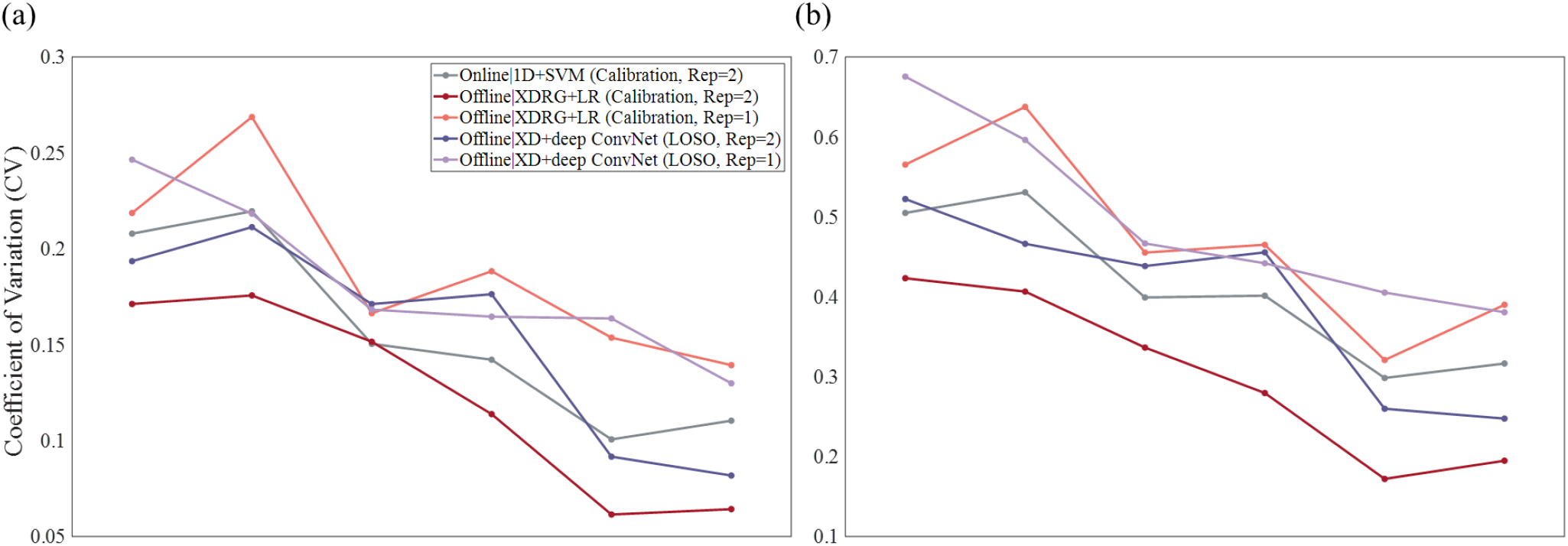
Individual variability of the performance of P300-Based BCIs measured by coefficient of variation (CV). The CV is measured across N=37 participants for six different P300-based BCI paradigms, including color-changing, icon-rotating and finger tapping stimulus designs combined with the counting and motor imagery (MI) tasks. The relationship of the CV and BCI paradigms is assessed for five different BCI setups: 1) online, 1D ERP feature vector, linear SVM classifier, individual calibration and 2 stimulus repetitions (Rep=2); 2) offline, xDAWN spatial filtering and Riemannian geometry transformation, logistic regression (LR) classifier, individual calibration and 2 stimulus repetitions (Rep=2); 3) offline, xDAWN spatial filtering and Riemannian geometry transformation, logistic regression (LR) classifier, individual calibration and single-trial (Rep=1); 4) offline, xDAWN spatial filtering, deepConvNet classifier, zero-calibration and 2 stimulus repetitions (Rep=2); and 5) offline, xDAWN spatial filtering, deepConvNet classifier, zero-calibration and single-trial (Rep=1). (a) CV in accuracy. (b) CV in ITR.

## 4. Discussion

In this study, we proposed a novel stimulus design employing the fingter-tapping animation relevant to target selection in the oddball task for P300-based BCIs. By eliciting more vivid ERPs using the proposed stimulus, aimed to address key challenges in P300-based BCIs, including repeated stimulus presentations, individual calibration, and variations in individual performance. P300-based BCIs with the finger-tapping stimlus showed the superior performance to those with conventional stimuli, reaching online accuracy of 91.17% across 37 participants to select one of the four commands for controlling an external device. Further offline optimization of computational models improved the accuracy of P300-based BCIs with the finger-tapping stimulus up to 97.3%. An offline test of single-trial P300-based BCIs revealed ITR of 59.91 bits/min while maintaining accuracy above 90%. Another offline across-subject evaluation demonstrated the plausibility of zero-calibration showing no difference in accuracy from individual calibration. Finally, using the finger-tapping stimulus further reduced variations in individual BCI performance compared to using conventional stimuli. Notably, greater performance enhancement by the finger-tapping stimulus in participants who showed relatively lower performance using conventional stimuli indicates that the proposed stimulus design was particularly effective for poorer BCI performers. Our results demonstrated the potential of our novel stimulus design to enhance BCI performance and realize plug-and-play BCI systems.

A key hypothesis driving our research was that elevated attention by an intuitive stimulus design could enhance ERPs leading to the improvement of P300-based BCIs. This was substantiated in our online experiment, where a mere alteration in the stimulus paradigm resulted in remarkable performance gains. Our experiment results demonstrate a crucial role of stimulus design in the development of P300-based BCIs, showcasing that eliciting reliable ERPs by well-designed stimuli would be as important as applying advanced computational algorithms to decode ERPs. This significant performance improvement, achieved with minimal stimulus repetitions, may pave the way for the development of high-performance BCIs more efficiently.

The optimization of computational models for ERPs elicited by the finger-tapping stimuli during the offline analysis further enhanced P300-based BCIs. Even with single-trial configurations, accuracy improved to 0.92 with an ITR nearing 60 bits/min. The accuracy achieved through individual calibration (97.3%) ranks the second-highest (99% being the hightest) among benchmarks in previous studies (see Table 4). However, note that the the study reporting the highest accuracy at 99% involved only two participants [48], potentially limiting its reliability. Moreover, the performance of single-trial BCIs built in this study surpasses existing benchmarks (Table 4), notably recording the highest ITR, to the best of our knowledge. These outcomes collectively suggest that our innovative stimulus paradigm enables us to build one of the most proficient P300-based BCIs to date.

**Table 4.** Comparative Analysis of Current Study with Previous Research in P300-Based BCIs

| Studies | Accuracy (%) | ITR (bits/min) | Number of Participants | Repetition of stimuli |
| --- | --- | --- | --- | --- |
| Present Study with Individual Calibration | 91.71 | 59.92 | 37 | Single-trial |
|  | 97.30 | 34.54 | 37 | 2 |
| Present Study with Zero Calibration | 87.75 | 52.61 | 37 | Single-trial |
| Kshirsagar et al., 2019 [48] | 92.64 | 55.45 | 9 | 15 |
| Kundu et al., 2020 [49] | 99 | - | 2 | 15 |
| Aygün et al., 2022 [50] | 94.56 | 7.73 | 30 | 15 |
| Blanco-Díaz et al., 2023 [51] | 76 | 11.38 | 10 | Single-trial |
| Du et al., 2023 [52] | 72.32 | - | 10 | Single-trial |

The outcomes of our optimization processes reveal some insights into the design of computational models for P300-based BCIs. First of all, spatial filgering such as xDAWN appears to be a key process for P300-based BCIs as shown in different analyses for single-trial BCIs or zero-calibration. Emplying more sophisticated methods such as RG transformation and deep ConvNet also contributed to improving performance. Yet, using both RG and deep neural networks did not improve performance further. It may imply that simply mixing different sophisticated algorithms would not help much for P300-based BCIs but a combination of different algorithms optimized for given ERP data would be more important.

High performance of zero-calibration P300-based BCIs demonstrated in this study may suggest that transfer learning without extensive data augmentation or domain adaptation is plausible for P300-based BCIs. It also points to the effectiveness of the xDAWN process and Riemannian geometry approach in trasfer learning as shown by previous reports [14]. But we also suspect that ERPs elicited by the finger-tapping stimuli would be more common among participants than those by conventional stimuli, supported by performance differences between stimulus designs in Table S2. Considering the universality of P300 components among people, elevating attention to a target stimulus by the proposed design would effectively mitigate variations of ERPs across participants.

Our results indicating better performance with finger-tapping than icon-rotating suggest that perceived task relevance likely plays a significant role in eliciting robust neural responses [53, 54]. Yet, there can be alternative explanations for the performance enhancement observed with the finger-tapping stimulus. One perspective is that the finger-tapping stimulus, composed of a more colorful image, could simply increase stimulus saliency. This increased saliency might lead to heightened bottom-up attention, consequently improving BCI performance, with minimal engagement of the task-relevance aspect of attention. Alternatively, the difference in performance between icon-rotating and finger tapping stimuli could be attributed to the representation of body-related stimuli. The animation of finger-tapping might draw more attention due to its relevance to bodily movements. Given that human perception tends to be more sensitive to stimuli related to human movements compared to other moving stimuli [55, 56], dynamic stimuli involving finger movements might attract greater attention than dynamic stimuli featuring rotating icons.

We observed no significant differences in BCI performance between the counting and MI tasks. This result may be attributed to the fact that participants needed to count a target only twice in each experimental block, alleviating the role of counting on engaging participants in the oddbal task. Also, the MI task might not be as effective as we had expected to increase attention as it depends heavily on the user’s ability to create vivid mental imagery. In fact, post-experiment feedback from some participants highlighted difficulties in performing MI along with the presented finger-tapping animation. Furthermore, the occurrence of movement-related cortical potentials (MRCPs) in the sensorimotor area [57] could potentially interfere with ERPs elicited by target stimuli. Nonetheless, the consistent high performance observed with the finger-tapping stimulus across different mental tasks suggests the feasibility of building P300-based BCIs without requiring a specific accompanying mental task, thereby simpifying BCI use with minimal cognitive effort.

Despite remarkable improvement of P300-based BCIs with innovative stimulus design, the present study has several limitations that need to be addressed further. As mentioned earlier, it is challenging to precisely determine why finger-tapping stimuli led to high performance. Understing a relationship between stimulus design and BCI performance in light of cognitive processing will be critical to advance the development of P300-based BCIs. Furthermore, the evaluation of BCI performance was limited to 15 blocks per test session, which may seem insufficient. However, this limitation was imposed by the experimental time constraints that require participants to maintain focus. Lastly, MI generally requires training and is subject to significant variability among participants, which may have limited its effectiveness in our study. Due to the rapid and repetitive nature of our paradigm, we were unable to develop a variety of MI tasks. Future research that includes a broader range of MI tasks may potentially lead to better performance.

In conclusion, our research marks a significant advancement in enhancing the performance of P300-based BCIs. Moving beyond conventional focus on computational models, this study emphasizes the importance of designing stimulus paradigms that align closely with human cognitive processes. By designing a user-centric stimulus based on attention mechanisms and cognitive engagement, we have laid a groundwork for more intuitive and efficient P300-based BCIs. These advancements may pave the way for the development of BCIs that are more accessible and user-friendly for everyday use. Particularly noteworthy is the potential demonstrated in this research for single-trial, zero-calibration P300-based BCIs, making a pivotal advancement. The next reserarch step naturally involves translating these results into highly usable plug-and-play BCI systems, aiming to broaden the spectrum of practical applications and make P300-based BCIs more readily available and convenient for diverse users. Our follow-up study will delve into the feasibility of such plug-and-ply P300-based BCIs.

## Acknowledgments

This work was supported by the National Research Foundation of Korea(NRF) grant funded by the Korea government(MSIT) (No. RS-2023-00302489).

