## Supplemental Table 1, Supplemental Table 2 for "Task-Relevant Stimulus Design Improves P300-Based Brain-Computer Interfaces"

**Table S1.** Individual calibration performance for all feature and classifier combinations. The 1D vector represents the concatenation of all channels by time, XD indicates the result of xDAWN filtering, and XDRG denotes the use of xDAWN filtering followed by Riemannian geometry. XDRG and EEGNet could not be implemented due to issues with input dimensions.

| Feature | | | 1D vector | | | | | | | | XD | | | | | | | | XDRG | | | | | | | |
|---|---|---|---|---|---|---|---|---|---|---|---|---|---|---|---|---|---|---|---|---|---|---|---|---|---|---|
| | | | Repetition = 1 | | | | Repetition = 2 | | | | Repetition = 1 | | | | Repetition = 2 | | | | Repetition = 1 | | | | Repetition = 2 | | | |
| Classification method | Stimuli | Mental task | Accuracy | | ITR(bits/min) | | Accuracy | | ITR(bits/min) | | Accuracy | | ITR(bits/min) | | Accuracy | | ITR(bits/min) | | Accuracy | | ITR(bits/min) | | Accuracy | | ITR(bits/min) | |
| | | | Mean | Std. | Mean | Std. | Mean | Std. | Mean | Std. | Mean | Std. | Mean | Std. | Mean | Std. | Mean | Std. | Mean | Std. | Mean | Std. | Mean | Std. | Mean | Std. |
| Linear SVM | Color change | Counting | 0.618 | 0.1768 | 20.3738 | 17.9888 | 0.7027 | 0.2076 | 15.2126 | 11.0513 | 0.6739 | 0.1497 | 24.4879 | 16.6409 | 0.7856 | 0.1657 | 19.5477 | 11.1768 | 0.7694 | 0.1697 | 37.3201 | 23.4405 | 0.8541 | 0.1538 | 24.6304 | 11.3528 |
| | | MI | 0.6396 | 0.1882 | 22.733 | 19.0507 | 0.7243 | 0.2109 | 16.7354 | 12.2263 | 0.755 | 0.184 | 35.6078 | 21.9929 | 0.7946 | 0.1738 | 20.2697 | 10.674 | 0.7405 | 0.2005 | 34.6771 | 22.7114 | 0.8739 | 0.1455 | 26.0554 | 10.7278 |
| | Icon-rotating | Counting | 0.6955 | 0.1645 | 27.2177 | 17.5382 | 0.7928 | 0.1546 | 19.6273 | 10.1148 | 0.7838 | 0.1328 | 36.6982 | 16.4186 | 0.836 | 0.1366 | 22.5508 | 10.1307 | 0.8378 | 0.1576 | 46.5993 | 22.7336 | 0.9093 | 0.1418 | 29.3468 | 10.5691 |
| | | MI | 0.6955 | 0.1622 | 27.1543 | 16.7173 | 0.8 | 0.1296 | 19.492 | 8.6551 | 0.7964 | 0.1396 | 38.8167 | 17.5073 | 0.8631 | 0.1515 | 25.2041 | 10.9018 | 0.836 | 0.166 | 46.9149 | 23.4888 | 0.9297 | 0.1165 | 30.805 | 9.3136 |
| | Finger-tapping | Counting | 0.7081 | 0.1628 | 28.5275 | 17.5216 | 0.8324 | 0.1273 | 22.1113 | 9.793 | 0.8396 | 0.1242 | 45.0564 | 18.7577 | 0.9027 | 0.1129 | 28.019 | 9.6725 | 0.9009 | 0.1428 | 56.8152 | 19.8254 | 0.9658 | 0.0855 | 34.0711 | 6.9436 |
| | | MI | 0.7261 | 0.1631 | 31.265 | 21.2133 | 0.8342 | 0.1253 | 22.0383 | 8.9538 | 0.8234 | 0.1229 | 42.4483 | 18.2743 | 0.9027 | 0.1061 | 27.5919 | 8.3588 | 0.8721 | 0.1251 | 51.09 | 20.91 | 0.9405 | 0.0699 | 30.8851 | 7.0484 |
| Logistic regression | Color change | Counting | 0.7117 | 0.1633 | 29.037 | 17.4363 | 0.7982 | 0.1637 | 20.1842 | 10.2314 | 0.6829 | 0.1606 | 25.9947 | 18.3656 | 0.7784 | 0.1626 | 18.8699 | 10.6256 | 0.791 | 0.173 | 40.2194 | 22.7363 | 0.8811 | 0.1508 | 27.0074 | 11.4289 |
| | | MI | 0.7441 | 0.1725 | 33.4848 | 20.2994 | 0.8108 | 0.1718 | 21.4348 | 11.1856 | 0.7405 | 0.1968 | 34.3144 | 22.0777 | 0.8 | 0.1707 | 20.6193 | 10.8423 | 0.7532 | 0.2024 | 36.5159 | 23.2801 | 0.8829 | 0.1551 | 27.139 | 11.0308 |
| | Icon-rotating | Counting | 0.782 | 0.1463 | 37.0221 | 16.6037 | 0.8595 | 0.135 | 24.3424 | 9.7472 | 0.7838 | 0.13 | 36.5927 | 16.4303 | 0.8324 | 0.1339 | 22.1352 | 9.6719 | 0.8595 | 0.143 | 49.8504 | 22.6944 | 0.9207 | 0.1395 | 30.437 | 10.2383 |
| | | MI | 0.7766 | 0.1687 | 37.4862 | 20.1719 | 0.8631 | 0.1154 | 24.3164 | 9.2492 | 0.8 | 0.1466 | 39.8686 | 19.2186 | 0.8793 | 0.1395 | 26.4397 | 10.7333 | 0.8631 | 0.1625 | 51.4776 | 23.9363 | 0.9315 | 0.106 | 30.6494 | 8.5665 |
| | Finger-tapping | Counting | 0.8306 | 0.133 | 44.3367 | 20.5678 | 0.9153 | 0.0856 | 28.5472 | 7.9913 | 0.8468 | 0.1304 | 46.6997 | 19.9397 | 0.9099 | 0.108 | 28.4723 | 8.9504 | 0.9171 | 0.1409 | 59.9156 | 19.221 | 0.973 | 0.0598 | 34.5455 | 5.9422 |
| | | MI | 0.7892 | 0.1536 | 38.8496 | 20.9475 | 0.8955 | 0.1049 | 26.9969 | 8.8398 | 0.8342 | 0.1203 | 44.0999 | 18.5514 | 0.8991 | 0.1118 | 27.4297 | 8.933 | 0.8775 | 0.1222 | 51.7459 | 20.1793 | 0.9586 | 0.0616 | 32.7974 | 6.389 |
| shallow ConvNet | Color change | Counting | 0.6306 | 0.2046 | 22.422 | 16.9046 | 0.7802 | 0.1865 | 19.4902 | 10.9877 | 0.6559 | 0.1863 | 24.1076 | 17.4007 | 0.7892 | 0.1882 | 20.2599 | 11.2704 | 0.3099 | 0.1407 | 3.0539 | 3.0801 | 0.4126 | 0.1811 | 3.5628 | 4.5945 |
| | | MI | 0.6432 | 0.2127 | 24.1958 | 20.1301 | 0.7766 | 0.1694 | 18.9692 | 11.1085 | 0.7063 | 0.1895 | 29.9025 | 21.0789 | 0.8252 | 0.1502 | 21.9727 | 10.3528 | 0.373 | 0.1591 | 4.959 | 7.7374 | 0.427 | 0.1674 | 3.5957 | 4.5292 |
| | Icon-rotating | Counting | 0.7477 | 0.1772 | 34.3201 | 21.2644 | 0.8486 | 0.1385 | 23.6618 | 10.2168 | 0.782 | 0.1529 | 37.5366 | 19.0191 | 0.8432 | 0.1269 | 23.0719 | 10.2462 | 0.3928 | 0.1639 | 5.7365 | 8.415 | 0.4216 | 0.1785 | 3.6835 | 5.0764 |
| | | MI | 0.7171 | 0.2116 | 32.2388 | 21.9428 | 0.8306 | 0.1836 | 23.4178 | 12.0537 | 0.7802 | 0.1624 | 37.8964 | 20.7059 | 0.8414 | 0.1666 | 23.8244 | 11.3491 | 0.3369 | 0.1594 | 4.1498 | 5.4237 | 0.4505 | 0.1688 | 4.1402 | 4.8275 |
| | Finger-tapping | Counting | 0.7676 | 0.1935 | 38.0306 | 23.8834 | 0.8865 | 0.1662 | 27.6423 | 10.638 | 0.8468 | 0.1294 | 46.6472 | 20.0713 | 0.9315 | 0.0923 | 30.4106 | 8.3195 | 0.382 | 0.1844 | 6.3073 | 11.2963 | 0.5135 | 0.1993 | 6.554 | 6.9979 |
| | | MI | 0.7874 | 0.1811 | 39.5366 | 20.9724 | 0.8955 | 0.1445 | 28.0481 | 10.6495 | 0.8378 | 0.1487 | 46.2701 | 22.0662 | 0.9027 | 0.1223 | 28.243 | 10.0368 | 0.4216 | 0.1879 | 7.89 | 12.0269 | 0.5514 | 0.2112 | 8.2539 | 9.328 |
| deep ConvNet | Color change | Counting | 0.7622 | 0.1802 | 36.1253 | 20.6641 | 0.8523 | 0.1508 | 24.1804 | 10.3554 | 0.7315 | 0.1703 | 31.9843 | 20.2868 | 0.8072 | 0.1562 | 20.8995 | 11.0702 | 0.5477 | 0.1951 | 15.3555 | 15.2428 | 0.6649 | 0.1802 | 12.478 | 9.7175 |
| | | MI | 0.7495 | 0.1835 | 35.0183 | 22.6604 | 0.8541 | 0.1464 | 24.3926 | 10.8662 | 0.7676 | 0.156 | 35.9394 | 20.065 | 0.818 | 0.1719 | 22.1718 | 11.7302 | 0.5586 | 0.1574 | 14.4012 | 11.5894 | 0.6631 | 0.1962 | 12.7925 | 10.4218 |
| | Icon-rotating | Counting | 0.8757 | 0.1259 | 51.0831 | 18.4512 | 0.9189 | 0.1258 | 29.9602 | 9.9884 | 0.8126 | 0.1617 | 42.5314 | 20.9836 | 0.8667 | 0.1449 | 25.5114 | 11.1367 | 0.5658 | 0.1789 | 15.8114 | 13.9517 | 0.7063 | 0.1725 | 14.52 | 9.8339 |
| | | MI | 0.8414 | 0.1534 | 46.8057 | 21.3053 | 0.9387 | 0.0921 | 31.3123 | 8.5696 | 0.8144 | 0.1572 | 42.5753 | 21.2478 | 0.8559 | 0.1503 | 24.7396 | 11.3092 | 0.6126 | 0.2205 | 22.0282 | 21.4013 | 0.7225 | 0.1882 | 15.885 | 10.873 |
| | Finger-tapping | Counting | 0.8811 | 0.1433 | 52.8582 | 19.3044 | 0.9568 | 0.0878 | 33.0501 | 7.3363 | 0.8559 | 0.1696 | 50.3758 | 23.5621 | 0.9405 | 0.0858 | 31.2044 | 7.7622 | 0.6631 | 0.1749 | 24.3746 | 17.9819 | 0.8108 | 0.1725 | 21.3852 | 10.8628 |
| | | MI | 0.8739 | 0.1131 | 50.4944 | 18.8002 | 0.9243 | 0.0905 | 29.656 | 8.4217 | 0.8649 | 0.1149 | 48.7519 | 17.8229 | 0.9351 | 0.0762 | 30.5093 | 7.6825 | 0.6937 | 0.1862 | 28.3681 | 21.3204 | 0.7387 | 0.1895 | 16.974 | 11.3826 |
| EEGNet | Color change | Counting | 0.7369 | 0.1625 | 32.1892 | 19.7002 | 0.8162 | 0.1628 | 21.613 | 10.8652 | 0.7207 | 0.177 | 31.0155 | 20.8134 | 0.8036 | 0.1579 | 20.5633 | 10.5691 | | | | | | | | |
| | | MI | 0.7135 | 0.1845 | 30.277 | 19.937 | 0.8324 | 0.156 | 22.6999 | 10.6889 | 0.7712 | 0.1607 | 36.6386 | 20.6655 | 0.845 | 0.1626 | 24.0239 | 11.2417 | | | | | | | | |
| | Icon-rotating | Counting | 0.8252 | 0.1435 | 43.5276 | 19.3076 | 0.8775 | 0.1356 | 26.0645 | 10.1953 | 0.8072 | 0.1386 | 40.8967 | 20.3053 | 0.8649 | 0.1461 | 25.3851 | 11.1442 | | | | | | | | |
| | | MI | 0.8144 | 0.1458 | 41.8688 | 19.4427 | 0.8991 | 0.1273 | 27.953 | 10.0285 | 0.8288 | 0.1419 | 44.1722 | 20.2354 | 0.8721 | 0.1318 | 25.623 | 10.497 | | | | | | | | |
| | Finger-tapping | Counting | 0.8468 | 0.1539 | 47.546 | 20.3379 | 0.9423 | 0.1068 | 31.9534 | 8.6917 | 0.8721 | 0.1271 | 51.0248 | 20.5743 | 0.9495 | 0.0692 | 31.8431 | 6.7552 | | | | | | | | |
| | | MI | 0.8414 | 0.1299 | 45.5703 | 19.277 | 0.9135 | 0.1142 | 29.1382 | 9.6459 | 0.8793 | 0.1265 | 52.0617 | 19.8036 | 0.9369 | 0.0719 | 30.5202 | 7.1604 | | | | | | | | |

**Table S2.** Calibration-free performance for all feature and classifier combinations. The 1D vector represents the concatenation of all channels by time, XD indicates the result of xDAWN filtering, and XDRG denotes the use of xDAWN filtering followed by Riemannian geometry. XDRG and EEGNet could not be implemented due to issues with input dimensions.

| Feature | | | 1D vector | | | | | | | | XD | | | | | | | | XDRG | | | | | | | |
|---|---|---|---|---|---|---|---|---|---|---|---|---|---|---|---|---|---|---|---|---|---|---|---|---|---|---|
| | | | Repetition = 1 | | | | Repetition = 2 | | | | Repetition = 1 | | | | Repetition = 2 | | | | Repetition = 1 | | | | Repetition = 2 | | | |
| Classification method | Stimuli | Mental task | Accuracy | | ITR(bits/min) | | Accuracy | | ITR(bits/min) | | Accuracy | | ITR(bits/min) | | Accuracy | | ITR(bits/min) | | Accuracy | | ITR(bits/min) | | Accuracy | | ITR(bits/min) | |
| | | | Mean | Std. | Mean | Std. | Mean | Std. | Mean | Std. | Mean | Std. | Mean | Std. | Mean | Std. | Mean | Std. | Mean | Std. | Mean | Std. | Mean | Std. | Mean | Std. |
| Linear SVM | Color change | Counting | 0.618 | 0.1768 | 20.3738 | 17.9888 | 0.7027 | 0.2076 | 15.2126 | 11.0513 | 0.6739 | 0.1497 | 24.4879 | 16.6409 | 0.7856 | 0.1657 | 19.5477 | 11.1768 | 0.4991 | 0.1711 | 10.9954 | 11.844 | 0.6468 | 0.1955 | 11.8924 | 9.9013 |
| | | MI | 0.6396 | 0.1882 | 22.733 | 19.0507 | 0.7243 | 0.2109 | 16.7354 | 12.2263 | 0.755 | 0.184 | 35.6078 | 21.9929 | 0.7946 | 0.1738 | 20.2697 | 10.674 | 0.5297 | 0.136 | 11.6673 | 9.3477 | 0.636 | 0.189 | 11.1363 | 8.3324 |
| | Icon-rotating | Counting | 0.6955 | 0.1645 | 27.2177 | 17.5382 | 0.7928 | 0.1546 | 19.6273 | 10.1148 | 0.7838 | 0.1328 | 36.6982 | 16.4186 | 0.836 | 0.1366 | 22.5508 | 10.1307 | 0.555 | 0.1499 | 13.8453 | 10.0122 | 0.7117 | 0.175 | 14.9939 | 10.6255 |
| | | MI | 0.6955 | 0.1622 | 27.1543 | 16.7173 | 0.8 | 0.1296 | 19.492 | 8.6551 | 0.7964 | 0.1396 | 38.8167 | 17.5073 | 0.8631 | 0.1515 | 25.2041 | 10.9018 | 0.6595 | 0.1646 | 23.2815 | 13.7469 | 0.7928 | 0.1447 | 19.3553 | 9.6431 |
| | Finger-tapping | Counting | 0.7081 | 0.1628 | 28.5275 | 17.5216 | 0.8324 | 0.1273 | 22.1113 | 9.793 | 0.8396 | 0.1242 | 45.0564 | 18.7577 | 0.9027 | 0.1129 | 28.019 | 9.6725 | 0.6667 | 0.1832 | 25.142 | 18.8374 | 0.8396 | 0.1105 | 22.377 | 9.2868 |
| | | MI | 0.7261 | 0.1631 | 31.265 | 21.2133 | 0.8342 | 0.1253 | 22.0383 | 8.9538 | 0.8234 | 0.1229 | 42.4483 | 18.2743 | 0.9027 | 0.1061 | 27.5919 | 8.3588 | 0.7207 | 0.1538 | 29.783 | 18.1047 | 0.8288 | 0.1419 | 22.1122 | 10.1488 |
| Logistic regression | Color change | Counting | 0.7117 | 0.1633 | 29.037 | 17.4363 | 0.7982 | 0.1637 | 20.1842 | 10.2314 | 0.6829 | 0.1606 | 25.9947 | 18.3656 | 0.7784 | 0.1626 | 18.8699 | 10.6256 | 0.6595 | 0.1438 | 22.6435 | 14.5942 | 0.7459 | 0.1624 | 16.4886 | 9.0955 |
| | | MI | 0.7441 | 0.1725 | 33.4848 | 20.2994 | 0.8108 | 0.1718 | 21.4348 | 11.1856 | 0.7405 | 0.1968 | 34.3144 | 22.0777 | 0.8 | 0.1707 | 20.6193 | 10.8423 | 0.7405 | 0.1624 | 32.536 | 18.9768 | 0.7892 | 0.1781 | 20.0891 | 11.191 |
| | Icon-rotating | Counting | 0.782 | 0.1463 | 37.0221 | 16.6037 | 0.8595 | 0.135 | 24.3424 | 9.7472 | 0.7838 | 0.13 | 36.5927 | 16.4303 | 0.8324 | 0.1339 | 22.1352 | 9.6719 | 0.7405 | 0.1585 | 32.3791 | 18.9159 | 0.8 | 0.1423 | 19.8899 | 9.804 |
| | | MI | 0.7766 | 0.1687 | 37.4862 | 20.1719 | 0.8631 | 0.1154 | 24.3164 | 9.2492 | 0.8 | 0.1466 | 39.8686 | 19.2186 | 0.8793 | 0.1395 | 26.4397 | 10.7333 | 0.7351 | 0.1575 | 31.6845 | 19.0707 | 0.8306 | 0.1761 | 23.0866 | 11.1589 |
| | Finger-tapping | Counting | 0.8306 | 0.133 | 44.3367 | 20.5678 | 0.9153 | 0.0856 | 28.5472 | 7.9913 | 0.8468 | 0.1304 | 46.6997 | 19.9397 | 0.9099 | 0.108 | 28.4723 | 8.9504 | 0.8378 | 0.1262 | 44.5753 | 17.7332 | 0.8937 | 0.1083 | 27.0338 | 9.4311 |
| | | MI | 0.7892 | 0.1536 | 38.8496 | 20.9475 | 0.8955 | 0.1049 | 26.9969 | 8.8398 | 0.8342 | 0.1203 | 44.0999 | 18.5514 | 0.8991 | 0.1118 | 27.4297 | 8.933 | 0.8234 | 0.1189 | 41.9902 | 16.4725 | 0.8721 | 0.1115 | 25.0863 | 9.342 |
| shallow ConvNet | Color change | Counting | 0.6306 | 0.2046 | 22.422 | 16.9046 | 0.7802 | 0.1865 | 19.4902 | 10.9877 | 0.6613 | 0.1681 | 23.6547 | 15.6228 | 0.7459 | 0.2066 | 17.8014 | 11.1398 | 0.4649 | 0.1486 | 8.2616 | 10.6994 | 0.5117 | 0.1626 | 5.7401 | 5.1679 |
| | | MI | 0.6432 | 0.2127 | 24.1958 | 20.1301 | 0.7766 | 0.1694 | 18.9692 | 11.1085 | 0.6883 | 0.1819 | 27.4604 | 19.858 | 0.7856 | 0.1847 | 20.1188 | 12.0692 | 0.4703 | 0.1539 | 8.7583 | 7.653 | 0.5874 | 0.1522 | 8.235 | 6.4802 |
| | Icon-rotating | Counting | 0.7477 | 0.1772 | 34.3201 | 21.2644 | 0.8486 | 0.1385 | 23.6618 | 10.2168 | 0.7405 | 0.1727 | 33.046 | 20.1456 | 0.8126 | 0.1464 | 20.9354 | 10.1666 | 0.4883 | 0.1743 | 10.4901 | 10.9728 | 0.6162 | 0.1636 | 9.6531 | 7.2479 |
| | | MI | 0.7171 | 0.2116 | 32.2388 | 21.9428 | 0.8306 | 0.1836 | 23.4178 | 12.0537 | 0.7694 | 0.156 | 35.9633 | 19.4438 | 0.8072 | 0.1705 | 21.1689 | 11.045 | 0.5586 | 0.1977 | 16.1646 | 13.8516 | 0.6072 | 0.1891 | 9.8024 | 7.6945 |
| | Finger-tapping | Counting | 0.7676 | 0.1935 | 38.0306 | 23.8834 | 0.8865 | 0.1662 | 27.6423 | 10.638 | 0.827 | 0.1374 | 43.701 | 19.9836 | 0.9207 | 0.09 | 29.3333 | 8.6167 | 0.6216 | 0.137 | 18.8466 | 12.6541 | 0.7027 | 0.1809 | 14.5188 | 9.9194 |
| | | MI | 0.7874 | 0.1811 | 39.5366 | 20.9724 | 0.8955 | 0.1445 | 28.0481 | 10.6495 | 0.827 | 0.1567 | 44.8506 | 22.0424 | 0.8937 | 0.1149 | 27.1439 | 9.5926 | 0.6036 | 0.1742 | 18.5929 | 13.9915 | 0.6937 | 0.214 | 14.9113 | 11.311 |
| deep ConvNet | Color change | Counting | 0.7622 | 0.1802 | 36.1253 | 20.6641 | 0.8523 | 0.1508 | 24.1804 | 10.3554 | 0.7081 | 0.1745 | 29.2397 | 19.7502 | 0.8054 | 0.1559 | 20.7008 | 10.8108 | 0.6414 | 0.1709 | 22.1356 | 17.545 | 0.7207 | 0.1669 | 15.0943 | 8.8607 |
| | | MI | 0.7495 | 0.1835 | 35.0183 | 22.6604 | 0.8541 | 0.1464 | 24.3926 | 10.8662 | 0.764 | 0.1667 | 36.0286 | 21.4802 | 0.8396 | 0.1774 | 23.7619 | 11.078 | 0.6937 | 0.1972 | 28.6396 | 19.6808 | 0.7766 | 0.1772 | 19.231 | 11.5675 |
| | Icon-rotating | Counting | 0.8757 | 0.1259 | 51.0831 | 18.4512 | 0.9189 | 0.1258 | 29.9602 | 9.9884 | 0.8054 | 0.1355 | 40.2737 | 18.7945 | 0.8685 | 0.1486 | 25.7929 | 11.3067 | 0.7045 | 0.132 | 26.896 | 14.2569 | 0.8234 | 0.1433 | 21.8505 | 10.7436 |
| | | MI | 0.8414 | 0.1534 | 46.8057 | 21.3053 | 0.9387 | 0.0921 | 31.3123 | 8.5696 | 0.8216 | 0.1352 | 42.5017 | 18.78 | 0.8613 | 0.1518 | 25.3105 | 11.5296 | 0.7315 | 0.1915 | 33.1201 | 22.14 | 0.8198 | 0.1562 | 21.553 | 9.92 |
| | Finger-tapping | Counting | 0.8811 | 0.1433 | 52.8582 | 19.3044 | 0.9568 | 0.0878 | 33.0501 | 7.3363 | 0.8775 | 0.1436 | 52.6084 | 21.316 | 0.9423 | 0.0863 | 31.5602 | 8.2012 | 0.8126 | 0.1395 | 41.5573 | 19.7235 | 0.8847 | 0.1073 | 26.0833 | 9.1024 |
| | | MI | 0.8739 | 0.1131 | 50.4944 | 18.8002 | 0.9243 | 0.0905 | 29.656 | 8.4217 | 0.8685 | 0.1128 | 49.5835 | 18.8692 | 0.9315 | 0.0762 | 30.0229 | 7.43 | 0.8198 | 0.1174 | 41.6471 | 17.6195 | 0.8721 | 0.1148 | 25.1111 | 9.3805 |
| EEGNet | Color change | Counting | 0.7369 | 0.1625 | 32.1892 | 19.7002 | 0.8162 | 0.1628 | 21.613 | 10.8652 | 0.7063 | 0.1876 | 29.8975 | 21.6046 | 0.7982 | 0.1629 | 20.3893 | 11.1168 | | | | | | | | |
| | | MI | 0.7135 | 0.1845 | 30.277 | 19.937 | 0.8324 | 0.156 | 22.6999 | 10.6889 | 0.7622 | 0.1544 | 34.9551 | 19.1445 | 0.8234 | 0.1672 | 22.4139 | 11.4874 | | | | | | | | |
| | Icon-rotating | Counting | 0.8252 | 0.1435 | 43.5276 | 19.3076 | 0.8775 | 0.1356 | 26.0645 | 10.1953 | 0.8054 | 0.1241 | 39.8372 | 18.2365 | 0.8468 | 0.1275 | 23.2347 | 9.8043 | | | | | | | | |
| | | MI | 0.8144 | 0.1458 | 41.8688 | 19.4427 | 0.8991 | 0.1273 | 27.953 | 10.0285 | 0.8252 | 0.1373 | 43.3105 | 19.5193 | 0.8523 | 0.1619 | 24.82 | 12.0133 | | | | | | | | |
| | Finger-tapping | Counting | 0.8468 | 0.1539 | 47.546 | 20.3379 | 0.9423 | 0.1068 | 31.9534 | 8.6917 | 0.8703 | 0.1143 | 49.6529 | 17.8794 | 0.9369 | 0.086 | 30.8102 | 7.8038 | | | | | | | | |
| | | MI | 0.8414 | 0.1299 | 45.5703 | 19.277 | 0.9135 | 0.1142 | 29.1382 | 9.6459 | 0.8703 | 0.1065 | 49.5315 | 17.9437 | 0.9207 | 0.0858 | 29.0068 | 7.7163 | | | | | | | | |
